# PEERS - an open science “Platform for the Exchange of Experimental Research Standards” in biomedicine

**DOI:** 10.1101/2021.07.31.454443

**Authors:** Annesha Sil, Anton Bespalov, Christina Dalla, Chantelle Ferland-Beckham, Arnoud Herremans, Konstantinos Karantzalos, Martien J. Kas, Nikolaos Kokras, Michael J. Parnham, Pavlina Pavlidi, Kostis Pristouris, Thomas Steckler, Gernot Riedel, Christoph H. Emmerich

## Abstract

Laboratory workflows and preclinical models have become increasingly diverse and complex. Confronted with the dilemma of assessing a multitude of information with ambiguous relevance for their specific experiments, scientists run the risk of overlooking critical factors that can influence the planning, conduct and results of studies and that should have been considered *a priori*. Negligence of such crucial information may result in sub-optimal study design and study execution, bringing into question the validity of generated outcomes. As a corollary, a lot of resources are wasted on biomedical research that turns out to be irreproducible and not sufficiently robust for further project development.

To address this problem, we present ‘PEERS’ (<u>P</u>latform for the <u>E</u>xchange of <u>E</u>xperimental <u>R</u>esearch <u>S</u>tandards), an open-access online platform that is built to aid scientists in determining which experimental factors and variables are most likely to affect the outcome of a specific test, model or assay and therefore ought to be considered during the design, execution and reporting stages.

The PEERS database is categorized into *in vivo* and *in vitro* experiments and provides lists of factors derived from scientific literature that have been deemed critical for experimentation. Most importantly, the platform is based on a structured and transparent system for rating the strength of evidence related to each identified factor and its relevance for a specific method/model. In this context, the rating procedure will not solely be limited to the PEERS working group but will also allow for a community-based grading of evidence.

To generate a proof-of-concept that the PEERS approach is feasible, we focused on a set of *in vitro* and *in vivo* methods from the neuroscience field, which are presented in this article. On the basis of the Open Field paradigm in rodents, we describe the selection of factors specific to each experimental setup and the rating system, but also discuss the identification of additional general items that transcend categories and individual tests. Moreover, we present a working format of the PEERS prototype with its structured information framework for embedding data and critical back end/front end user functionalities. Here, PEERS not only offers users the possibility to search for information to facilitate experimental rigor, but also draws on the engagement of the scientific community to actively expand the information contained within the platform through a standardized approach to data curation and knowledge engineering.

As the database grows and benefits become more apparent, we will expand the scope of PEERS to any area of applied biomedical research.

Collectively, by helping scientists to search for specific factors relevant to their experiments, and to share experimental knowledge in a standardized manner, PEERS will serve as the ultimate exchange and analysis tool to enhance data validity and robustness as well as the reproducibility of preclinical research. PEERS offers a vetted, independent tool by which to judge the quality of information available on a certain test or model, identifies knowledge gaps and provides guidance on the key methodological considerations that should be prioritized to ensure that preclinical research is conducted to the highest standards and best practice.

## 1. Introduction and rationale

Biomedical research, particularly in the preclinical sphere, has been subject to scrutiny for the low levels of reproducibility that continue to persist across laboratories (Ioannidis, 2005). Reproducibility in this context refers to the ability to corroborate results of a previous study by conducting new experiments with the same experimental design but collecting new and independent data sets. Reproducibility checks are common in fields like physics (CERN Education, Communications and Outreach Group, 2018), but rarer in biological disciplines such as neuroscience and pharmacotherapy, which are increasingly facing a ‘reproducibility crisis’ (Bespalov et al., 2016; Bespalov and Steckler, 2018; Botvinik-Nezer et al., 2020). Even though a high risk of failure to repeat experiments between laboratories is an inherent part of developing innovative therapies, some risks can be greatly reduced and avoided by adherence to evidence-based research practices using clearly identified measures to improve research rigor (Vollert et al., 2020; Bespalov et al., 2021; Emmerich et al., 2021). Alternative initiatives have been introduced to increase data reporting and harmonization across laboratories [ARRIVE 2.0 (Percie du Sert et al., 2020); EQUATOR network (Simera, 2008); The International Brain Laboratory (International Brain Laboratory et al., 2017); FAIRsharing Information Resource (Sansone et al., 2019)], improve data management and analysis [(Pistoia Alliance Database (Makarov et al., 2021); NINDS Common Data Elements (Stone, 2010); FITBIR: Traumatic Brain Injury network (Tosetti et al., 2013); FITBIR: Preclinical Traumatic Brain Injury Common Data Elements (LaPlaca et al., 2021)], or publish novel methods and their refinements (Norecopa; Current Protocols in Neuroscience; protocols.io; The Journal of Neuroscience Methods). However, extrinsic and intrinsic factors that affect study outcomes in biomedical research have not yet been systematically considered or weighted and are the subject of ‘PEERS’ (Platform for the Exchange of Experimental Research Standards). This makes PEERS a unique addition to this eclectic list of well-established resources.

The rationale for PEERS is as follows: Laboratory workflows and preclinical models have become increasingly diverse and complex. Although the mechanics of many experimental paradigms are well explored and usually repeatable across laboratories and even across national/continental boundaries (Robinson et al., 2018; Aguillon-Rodriguez et al., 2021), data can be highly variable and are often inconsistent. Multiple attempts have been made to overcome this issue, but even efforts in which experimental conditions were fully standardized between laboratories have not been completely successful. This may not be surprising given that behavioral testing, for example, is sensitive to environmental factors such as housing conditions (background noise, olfactory cues), experimenter interactions, sex or the strain under investigation (Sousa et al., 2006; Bohlen et al., 2014; Riedel et al., 2018; Pawluski et al., 2020; Butlen-Ducuing et al., 2021). Many multi-laboratory studies have also observed significant differences between mouse strains and interactions of genotype × laboratory despite efforts to rigorously standardize both housing conditions and experimental design (Wolfer et al., 2004; Richter et al., 2011). Taking together, there are many variables/factors that can affect an experiment and the outcome of a study.

A proper catalogue of these influencing factors, including the scientific evidence combined with a rating of its strength, is missing to date and PEERS seeks to fill this gap and aims to guide scientists by advising which factors need to be monitored, recorded or reported. Figure 1 represents the overarching concept of the PEERS platform (Fig. 1).

**Figure 1:**
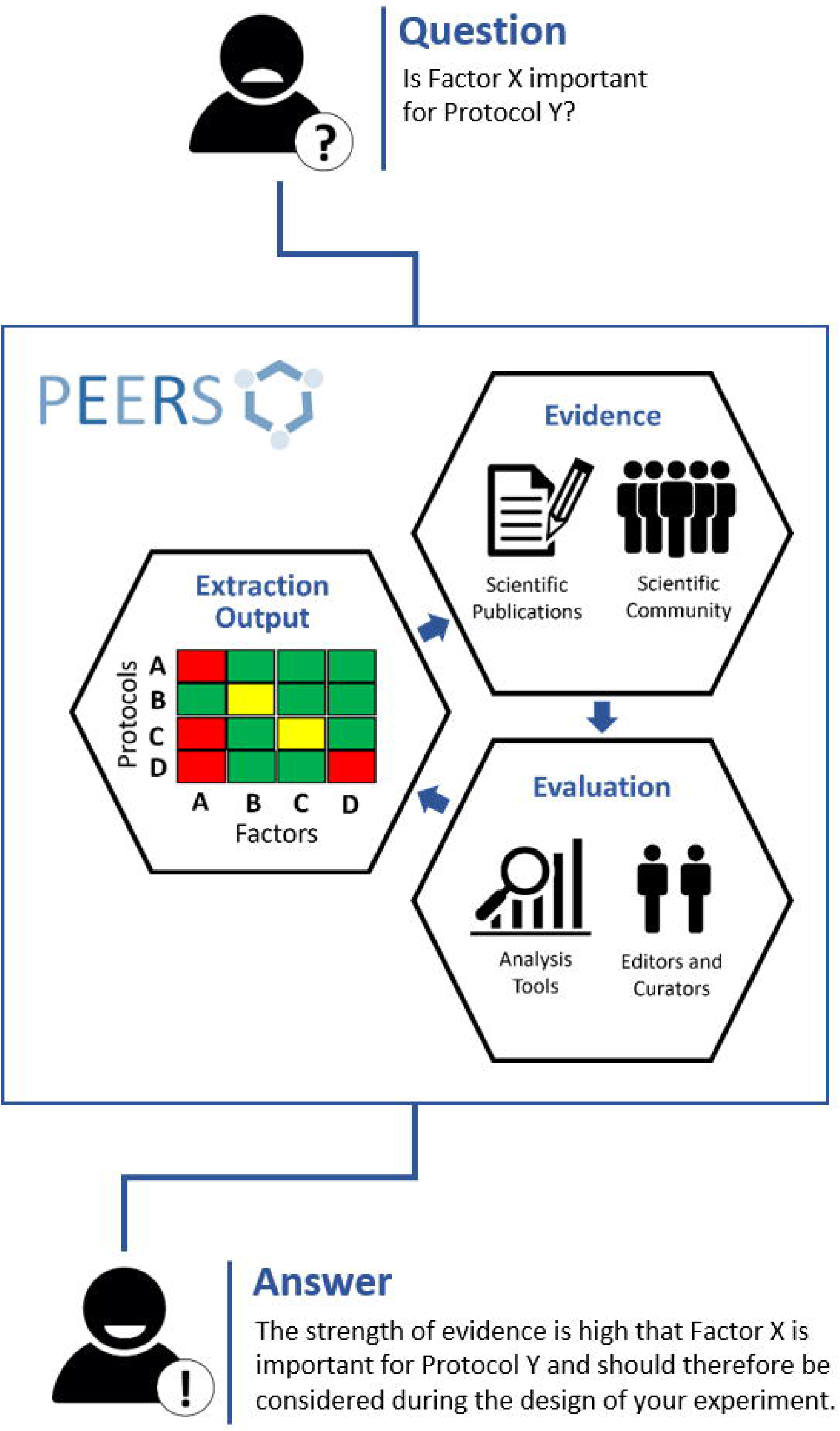
Outline of the PEERS concept and workflow (the 3Es). To understand whether specific factors are relevant for certain methods/models (‘protocols’), the PEERS workflow is based on different steps to a) collect information about selected factors/protocols from publications or the scientific community (‘**Evidence**’); b) rate the strength of this information and provide mechanisms for editing, curating and maintaining the information/database (‘**Evaluation**’); and c) present the outcome in a user-friendly and digestible form (‘**Extraction** Output’) so that users will be provided with an answer helpful for their planned experiments.

## 2. The PEERS Solution

To mitigate some of the above issues, we have developed PEERS, an open-access online platform that seeks to aid scientists in determining which experimental factors (or variables) most likely affect the outcome of a specific test, model or assay and therefore deserve consideration *prior* to study design, execution and reporting. Our overarching ambition is to develop PEERS into a *one-stop exchange and reporting tool* for extrinsic and intrinsic factors underlying variability in study outcomes and thereby undermining scientific progress. At the same time, PEERS offers a vetted, independent perspective by which the quality of information available on a certain test or model can be judged. It will also identify knowledge gaps and provide guidance on key methodological considerations that should be prioritized to ensure that preclinical research is conducted to the highest standards and incorporates best practice.

### 2.1. The PEERS Consortium

The PEERS project can be traced back to the Preclinical Data Forum (PDF) (https://www.preclinicaldataforum.org/), a network financially and organizationally supported by the European College of Neuropsychopharmacology (ECNP) and Cohen Veterans Bioscience (CVB). The PDF focuses on robustness, reproducibility, translatability and transparency of reporting preclinical data and consists of a multinational consortium of specialist researchers from research institutions, universities, pharma companies, SMEs and publishers. Originating from the PDF, the PEERS Working Group (AS, CD, CFB, AH, KK, MJK, NK, KP, GR, CHE) is currently funded by CVB during its initiation phase. The Working Group consists broadly of the ‘scientific arm’ with long-standing expertise in neuroscience, reproducibility and improvements in data quality across academic and industrial preclinical biomedical research. The ‘scientific arm’ of the group is complemented by the strong software and machine learning expertise of the ‘software arm’ which translates the scientific input provided into an easy to navigate open access platform.

Therefore, our initial focus is on *in vivo* and *in vitro* methods commonly utilized in neuroscience research. Since the inaugural meeting on 10 September 2020, the implementation of a principal concept was agreed, and partners have contributed to different work-packages.

### 2.2. How does PEERS work?

#### 2.2.1 The PEERS database and its front and back-end functionalities

Figure 2 represents the overall structure of the platform with the front and back-end functionalities represented. The front end contains a data input module which allows registered users to add either new methods/models (here termed ‘**protocols’**) or provide add-on information to existing protocols, but also the data search and the data extraction modules to be used by a typical user for the examination of databases. The back end of the PEERS database contains the processes to collect and analyze information related to the selected protocols. The relevance of specific factors for the outcome of these protocols is analyzed based on a detailed scoring system, representing a central element of the PEERS working prototype. The different steps involved in setting up this platform are discussed in the following sections by following the 3Es identified in Fig. 1.

**Figure 2:**
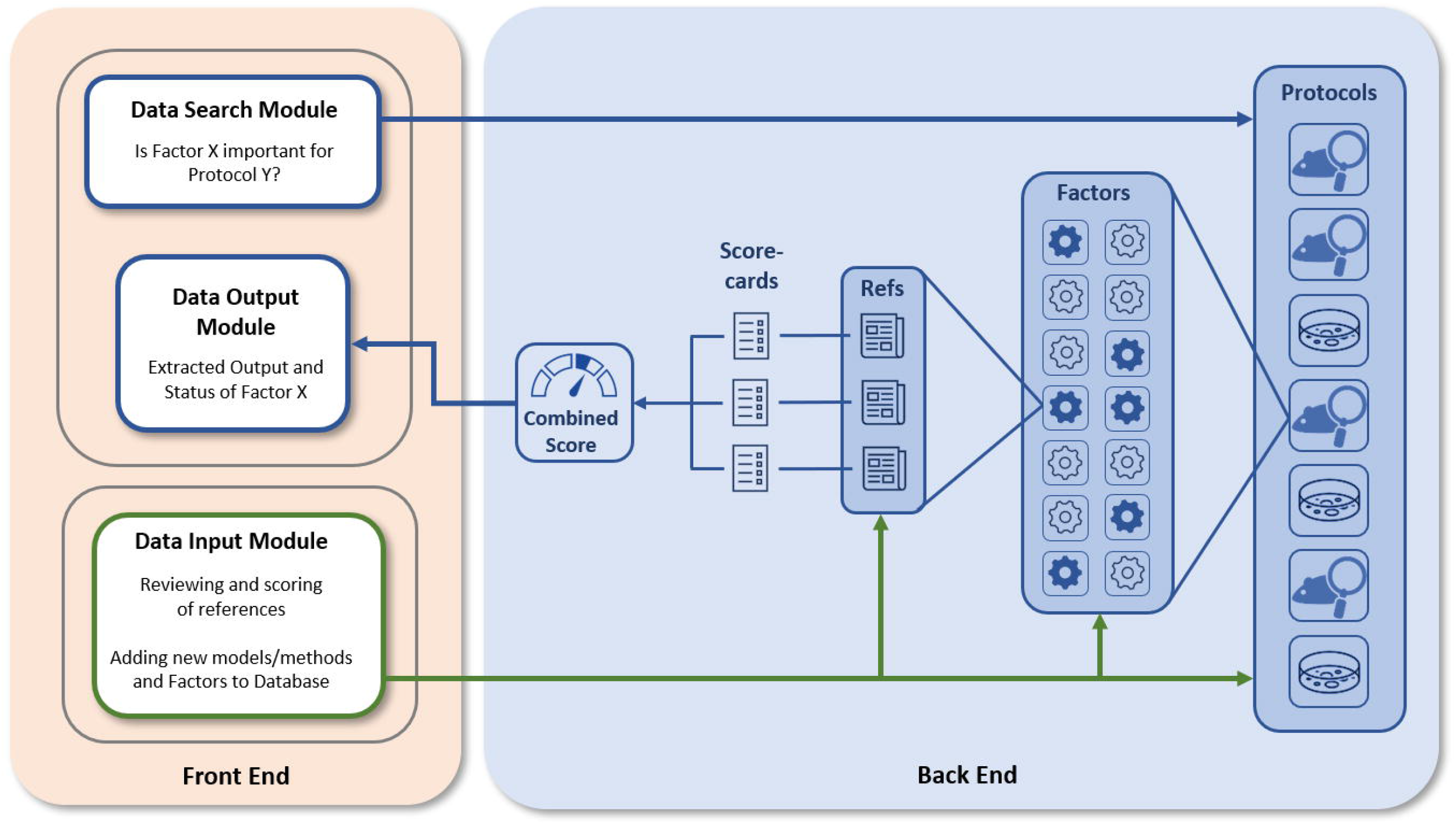
PEERS platform structure. Users can interact with the PEERS platform (blue arrows) by searching for or adding information (Front End Modules). The PEERS database (Back End) consists of various protocols, for which generic and specific factors and related references have been identified. The Quality of Evidence for the importance of certain factors is evaluated using scorecards and a summary is presented by visualizing results in the user interface. Users can contribute by adding new protocols or factors and by scoring relevant references (green arrows).

##### 2.2.1.1. Selection of *in vivo/in vitro* protocols

For the working prototype and as a proof-of-concept, four *in-vivo* and four *in vitro* protocols were identified, based on (1) how commonly they are used in neuroscience (and by extension the literature available on them), and (2) the expertise of the core group (see Table 1).

**Table 1:** The four initial *in vivo* and *in vitro* neuroscience protocols selected for the PEERS platform.

| IN VIVO | IN VITRO |
| --- | --- |
| Open Field | Western blotting |
| Water Maze | PCR |
| EEG | ELISA |
| Conditioned Place Preference | Calcium Imaging |

##### 2.2.1.2. Selection of ‘factors’

The central elements of PEERS are the ‘**factors**’, defined as any aspect of a study that i) can affect the study outcome, and therefore, ii) information to incorporate them into the study design (e.g., ignore/control, monitor, report them) is required. Factors were divided into two categories: 1) generic factors relevant to all protocols (e.g., strain of animals) and 2) specific factors relevant to and affecting specific protocols only (e.g., water temperature for water maze) by utilizing expert opinion from within the PEERS consortium. Users of the platform can search for these factors depending on the protocols and outcomes they are interested in. Representative tables of factors for one *in vivo* (Open Field test) and one *in vitro* method (Western blotting) can be found in the Supplemental section (Table S1 and S2).

##### 2.2.1.3. Collection of references that report on selected ‘factors’ – The Evidence

To identify and collate references that study the importance of a specific factor for a selected protocol and its outcome, an extensive review of published literature via the PubMed and EMBASE databases was conducted. These included references dealing with either a factor of interest that had an effect on the protocol outcome or one that had no effect on the outcome when manipulated. A representative table of factors with references for the Open Field protocol can be found in the Supplemental section (Table S1).

#### 2.2.2. Grading of Evidence: description of structured approach

Within the PEERS database, we provide references for each factor that has been scrutinized. We have gone one step further by providing a grading of the strength of this evidence (either positive or negative) so that examination of a specific factor in the database provides the user with an extracted summary of all relevant papers and their scores from one or more assessors (scorecards). This required the development of a generic ‘checklist’ to determine the quality of each paper the details of which are described in the following section.

##### 2.2.2.1. Checklist for grading of evidence/publications – The Evaluation

Concurrently with the identification of experimental factors and the review of literature, novel detailed ‘scorecards’ to evaluate the quality of scientific evidence were refined through multiple Delphi rounds within the PEERS Working Group. These contain a checklist with two main domains – *Methods* and *Results*. The elements of these domains were determined based on ARRIVE 2.0 ‘Essential 10’ and recommendations of the EQIPD consortium (Percie du Sert et al., 2020; Vollert et al., 2020). The *Methods* domain assesses the adherence to these guidelines with a maximum score of 10 (essentially one point for each of the 10 items – or fractions of 1 if items are only covered partially - or zero points if specific items are not covered at all). The *Results* domain meanwhile aims at evaluating the quality and suitability of the results and analyses, and again a score of 10 was awarded if all items were sufficiently addressed. The scorecards constitute a unique feature of the PEERS database because not only do they evaluate reporting of the methods in any paper but also take into account the suitability and strength of the results presented. Ideally, each reference is evaluated by two or more assessors to remove any source of bias. Table 2 depicts the checklist score utilized for the *in vivo* protocols.

**Table 2:**
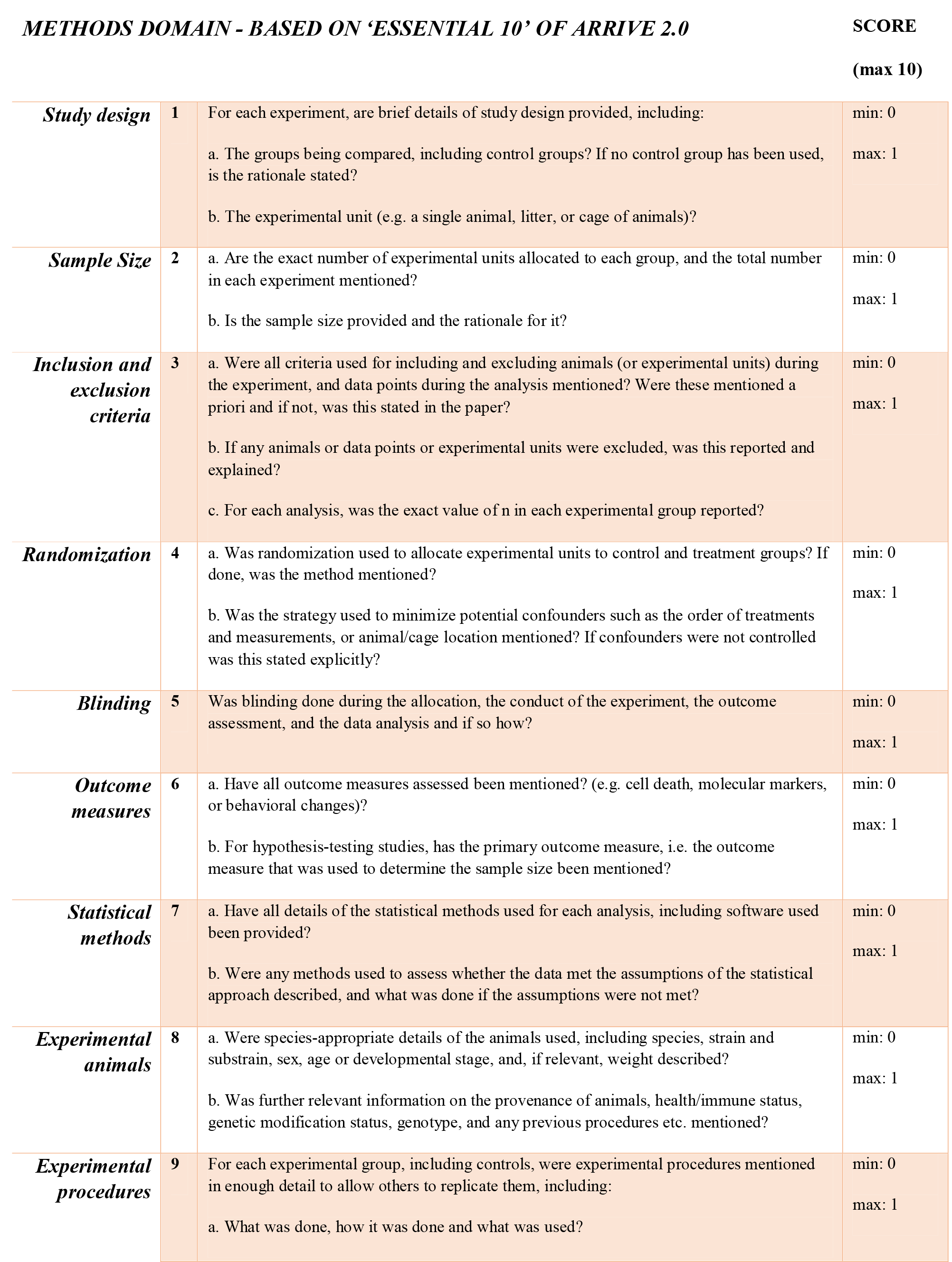
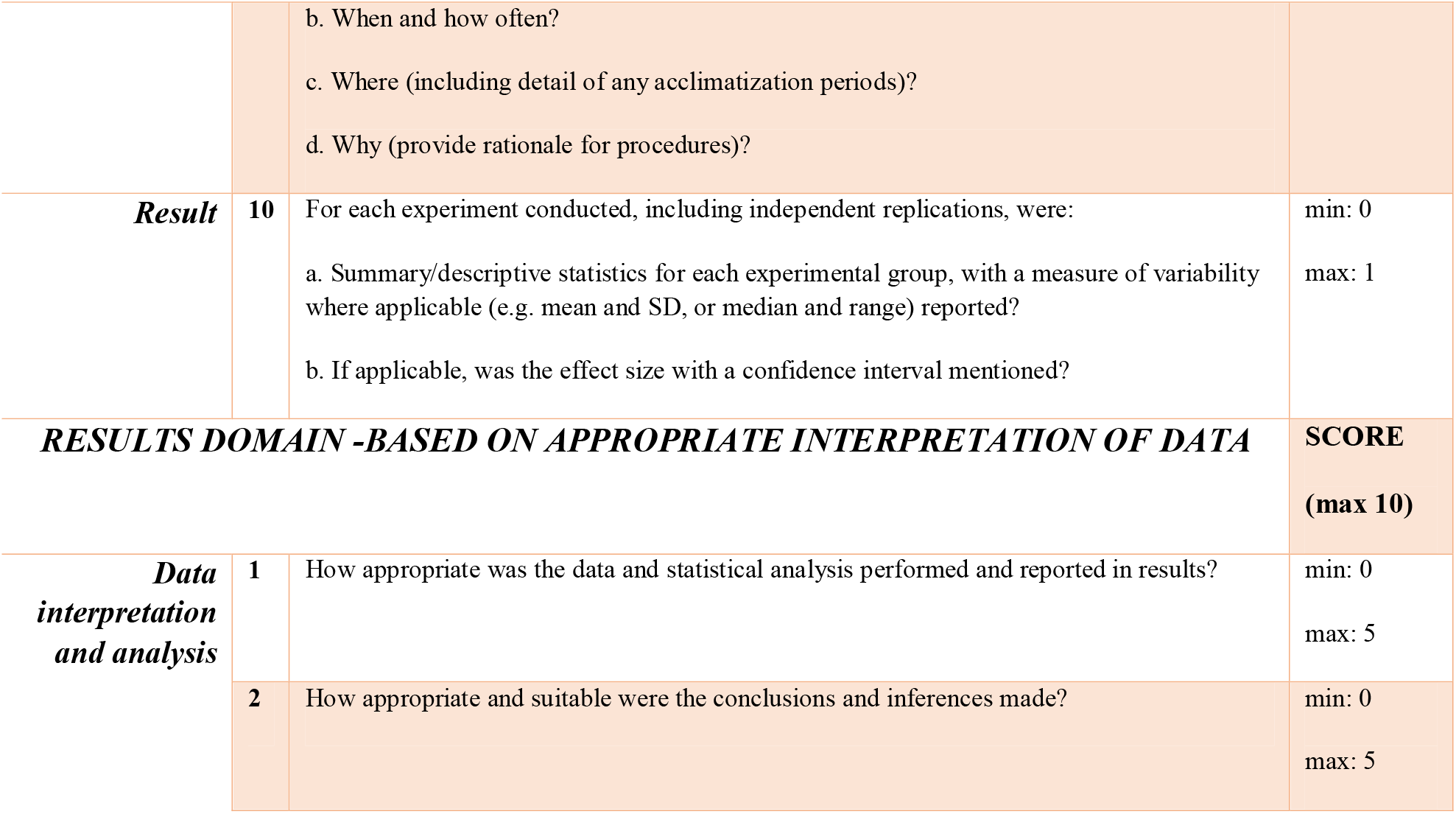
‘Scorecard’ containing checklist for the *in vivo* protocols.

**METHODS DOMAIN - BASED ON ‘ESSENTIAL 10’ OF ARRIVE 2.0** **SCORE** **(max 10)**
|  |  |  |  |
| --- | --- | --- | --- |
| **Study design** | **1** | For each experiment, are brief details of study design provided, including:<br>a. The groups being compared, including control groups? If no control group has been used, is the rationale stated?<br>b. The experimental unit (e.g. a single animal, litter, or cage of animals)? | min: 0<br>max: 1 |
| **Sample Size** | **2** | a. Are the exact number of experimental units allocated to each group, and the total number in each experiment mentioned?<br>b. Is the sample size provided and the rationale for it? | min: 0<br>max: 1 |
| **Inclusion and exclusion criteria** | **3** | a. Were all criteria used for including and excluding animals (or experimental units) during the experiment, and data points during the analysis mentioned? Were these mentioned a priori and if not, was this stated in the paper?<br>b. If any animals or data points or experimental units were excluded, was this reported and explained?<br>c. For each analysis, was the exact value of n in each experimental group reported? | min: 0<br>max: 1 |
| **Randomization** | **4** | a. Was randomization used to allocate experimental units to control and treatment groups? If done, was the method mentioned?<br>b. Was the strategy used to minimize potential confounders such as the order of treatments and measurements, or animal/cage location mentioned? If confounders were not controlled was this stated explicitly? | min: 0<br>max: 1 |
| **Blinding** | **5** | Was blinding done during the allocation, the conduct of the experiment, the outcome assessment, and the data analysis and if so how? | min: 0<br>max: 1 |
| **Outcome measures** | **6** | a. Have all outcome measures assessed been mentioned? (e.g. cell death, molecular markers, or behavioral changes)?<br>b. For hypothesis-testing studies, has the primary outcome measure, i.e. the outcome measure that was used to determine the sample size been mentioned? | min: 0<br>max: 1 |
| **Statistical methods** | **7** | a. Have all details of the statistical methods used for each analysis, including software used been provided?<br>b. Were any methods used to assess whether the data met the assumptions of the statistical approach described, and what was done if the assumptions were not met? | min: 0<br>max: 1 |
| **Experimental animals** | **8** | a. Were species-appropriate details of the animals used, including species, strain and substrain, sex, age or developmental stage, and, if relevant, weight described?<br>b. Was further relevant information on the provenance of animals, health/immune status, genetic modification status, genotype, and any previous procedures etc. mentioned? | min: 0<br>max: 1 |
| **Experimental procedures** | **9** | For each experimental group, including controls, were experimental procedures mentioned in enough detail to allow others to replicate them, including:<br>a. What was done, how it was done and what was used? | min: 0<br>max: 1 |
|  |  | b. When and how often?<br>c. Where (including detail of any acclimatization periods)?<br>d. Why (provide rationale for procedures)? |  |
| **Result** | **10** | For each experiment conducted, including independent replications, were:<br>a. Summary/descriptive statistics for each experimental group, with a measure of variability where applicable (e.g. mean and SD, or median and range) reported?<br>b. If applicable, was the effect size with a confidence interval mentioned? | min: 0<br>max: 1 |
| **RESULTS DOMAIN -BASED ON APPROPRIATE INTERPRETATION OF DATA** |  |  | **SCORE**<br>**(max 10)** |
| **Data interpretation and analysis** | **1** | How appropriate was the data and statistical analysis performed and reported in results? | min: 0<br>max: 5 |
|  | **2** | How appropriate and suitable were the conclusions and inferences made? | min: 0<br>max: 5 |

Multiple scorecards (from different assessors) dealing with the same factor/reference as well as the overarching score derived from the description of methods and results are freely accessible on the PEERS platform for detailed information.

##### 2.2.2.2. Overall grading of evidence – The Extraction

Following on from the individual scoring of each paper, an algorithm commonly used for meta-analyses (Neyeloff et al., 2012) that quantifies the degree of reviewer consensus (heterogeneity of grading) was used to provide an adjusted grade for the extracted output taking together all scores of all reviewers.

This grading system was then simplified to establish the overall scores for each factor into high (>14/20) / medium (5-13/20) / low (<5/20) quality (or no evidence), which is then displayed for users on the front-end of the platform.

#### 2.2.3. Platform users and contributors

All first-time users on the PEERS platform need to register and accept the PEERS Code of Conduct (CoC-see below in Section 3.1) in order to use PEERS. This includes a small profile page where users can fill in details of their present affiliation, level of expertise and areas of interest. This also allows the different users to be guided to areas of interest that match up with their profile. The platform will display the active users on the platform and will also show details of their involvement and contribution to different protocols (optional, only if desired). With time, these measurable contributions/metrics can be used by users to demonstrate their effort, time involvement and value.

Until a critical number of users is reached, the scoring of publications by new (and unexperienced) users will be moderated briefly by an experienced member of the PEERS Working Group. However, as mentioned in Section 2.2.2.1, the scorecard checklists are kept as simple and intuitive as possible (e.g. by adhering to the ARRIVE 2.0 guidelines for methods reporting) so that scoring of publications is neither time-consuming nor difficult.

Once registered, users can ask questions about the relevance of specific factors by interacting with the ‘Data Search Module’ (Fig. 2). Additionally, they can also contribute to the reviewing and scoring of references using the ‘Data Input Module’. Ultimately, PEERS extracts an output for the user summarizing the ‘status’ of the factor of interest and detailing the scientific strength available that the factor may indeed influence the design, conduct and reporting phase of the protocol of choice.

### 2.3. Description of the PEERS prototype

The current PEERS prototype consists of a web application such that any expert, after registration, may insert data and review any existing protocol datasets. By implementing a relational schema, provisions have been made for easy data transformation using semi-structured formats such as JSON and XML, so they are ready for sharing with other applications or systems through an Application Programming Interface (API). The central entities to be collected and stored in the PEERS database are the *in vivo* and *in vitro* protocols including all related factors, references, and scorecards. The prototype is set up using the popular ReactJS library with a simple and effective design provided by the Semantic UI framework, while data are managed and stored through CVB’s *BRAINS Commons* platform, a cloud-based platform for computational discovery across brain disease. The Minimum Viable Product (MVP) includes a user management, authentication, and authorization module so that the access/contributions of each user can be tracked and presented in the final dataset. This feature facilitates implementation of collaborative elements which PEERS seeks to integrate. The application will be accessible using any web browser via the following link: https://www.braincommons.org/peers-platform/.

### 2.4. An example: The ‘Open Field’ protocol

The following example demonstrates how the different back-end functionalities of PEERS will translate into the front-end ‘Extracted Output’ displayed on the PEERS platform when users aim to retrieve information about specific factors for the Open Field *in vivo* protocol (Fig. 3).

**Figure 3:**
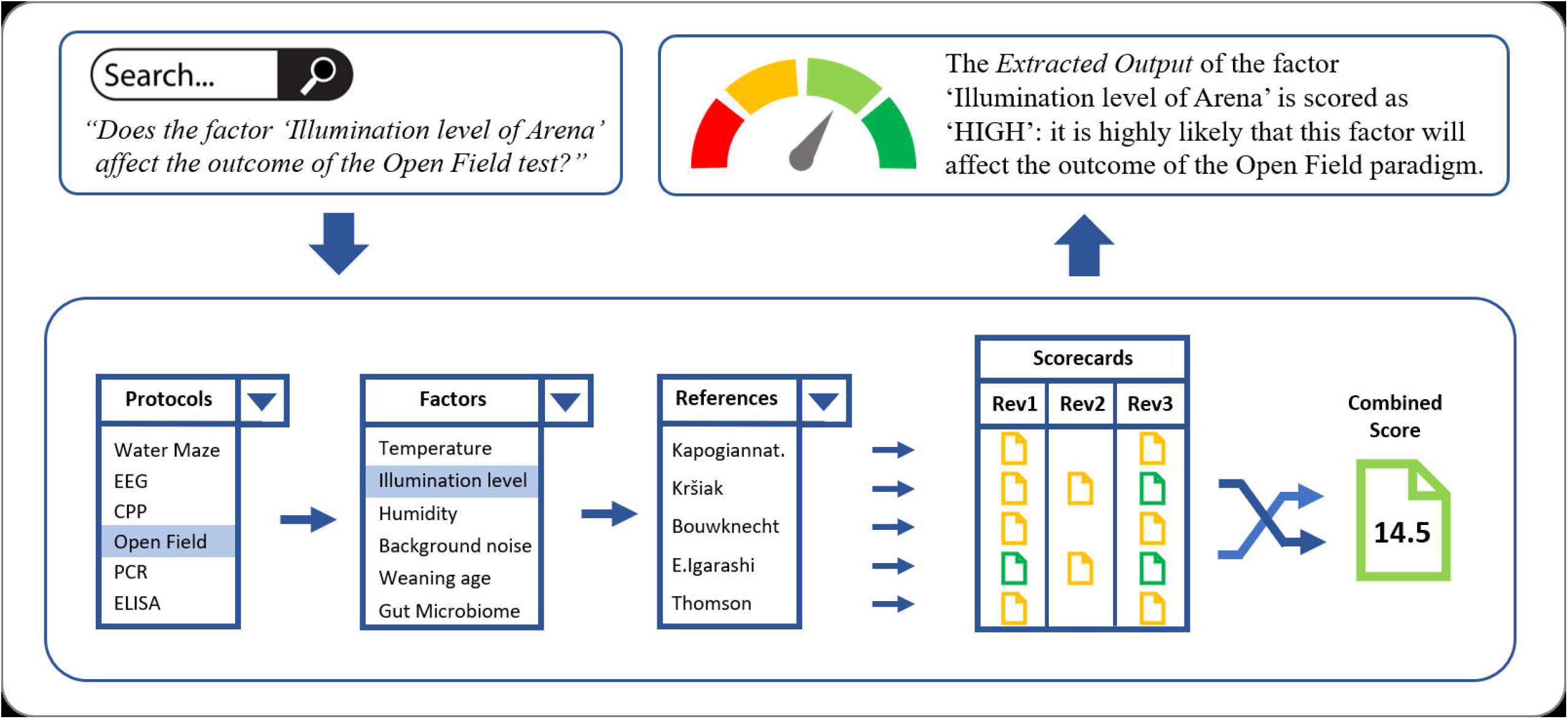
The ‘Open Field’ protocol example, demonstrating how the different back-end functionalities of PEERS will translate into the ‘Extracted Output’, presented to PEERS users. A search query for a specific factor/protocol will lead to the selection of all relevant references from the PEERS database dealing with the factor of interest (e.g., the ‘illumination level of the arena’). Based on the scorecards for these references the combined score is calculated which translates into the overall extracted output for the selected factor/protocol combination. This status is then presented to the user. Users also have access to all scorecards to understand how the overall grading of evidence was achieved.

Accessing the Open Field test for the first time, users of PEERS might want to ask the question: ‘*Do the factors ‘Sex’ (generic) and ‘Illumination level of arena’ (specific) affect the outcome of the Open Field test?’*

a. As a first step, users accessing the platform can input their query into the search module such as: Will the factor ‘Sex’ affect the outcome of an Open Field test? or Will the factor ‘Illumination level of arena’ affect the outcome of an Open Field test? and click *Search*. All factors and tests will be displayed in drop-down menus.
b. Subsequently, the PEERS database will i) locate all the factors pertaining to the Open Field protocol and will select ‘*Sex*’ or ‘*Illumination level of arena*’ from these, ii) generate the list of references (via DOIs) for the two factors, iii) provide scorecards scored by reviewers for each of the references, (iv) use the mathematical model to resolve any discrepancies between reviewers/multiple references, and (v) generate the overall extracted output of the evidence for ‘*Sex’* or ‘*Illumination level of Arena’* as either high (>14/20) / medium (5-13/20) / low (<5/20) quality (or no evidence).
c. The extracted output status of the two factors will then be visualized as ‘HIGH’ in this case - meaning that it is highly likely that both factors ‘Sex’ and ‘Illumination level of arena’ can affect the outcome of the Open Field paradigm. To ensure full transparency, users will also have access to all scorecards and related references for each of the papers scored and can follow each step to understand how the overall grading of evidence was achieved.

## 3. Future directions and outlook

### 3.1. Proposed curation mechanisms and community engagement

#### 3.1.1. Wiki-like functionality

To involve the scientific community at large in building and growing the PEERS database, a wiki-like functionality is adopted to allow the collaborative modification and addition of content and structure. The wiki concept ensures that all stakeholders (including early career researchers – PhD students; research assistants and fellows – and established professionals) can actively participate in the curation and reviewing process of evidence pertaining to a protocol using a standardized approach to evaluating the evidence. The presence of multiple reviewers for each factor and protocol will ensure that there is no bias while the meta-analysis approach described above will be utilized to resolve any disagreement between reviewers and adjust the strength of evidence should new information become available.

Editing, curating, and maintaining the PEERS platform is integral to the process and the Working Group is proposing several measures to credit any contributor to the review/editing/curating process. Some of these proposals include making PEERS recommendations citable like conventional publications or developing the platform such that contributors’ names appear on protocols they have contributed to. In addition to voluntary contributions, qualified staff will ensure maintenance, quality management of the content on the database, sustainability and project management functionalities where needed.

#### 3.1.2. Governance: The PEERS Code of Conduct

The PEERS platform aims to be a community-driven resource which will be curated and updated regularly in an open fashion. We expect biomedical researchers from both academia and industry all over the world to be members and contributors of this community. Above all, we would expect this community to be respectful and engaged so we can reach a broader audience and be helpful to scientists at different stages of their career and in different research environments. Therefore, we will implement a PEERS Code of Conduct (CoC) and all users will have to accept the PEERS CoC before becoming contributors or active users. The CoC will be formulated as a guide to make the community-driven nature of the platform productive and welcoming.

However, violations of the CoC will affect the user’s ability to contribute to the PEERS database and to score papers and engage with the wider PEERS community. Often users will be scoring the quality of methods and results presented in a scientific paper and these are bound to have consequences for other users, colleagues or authors of that paper and therefore, it is important to be respectful, fair and open. Users must not allow any personal prejudices or preferences to overshadow the scoring of any papers and must judge a paper purely on the content presented in it.

If any disagreements do arise, they should be dealt with in a mature fashion and when possible, informally. However, if the informal processes seem inadequate to resolve any conflicts, PEERS will establish a structured procedure to deal with any complaints or report against any problematic users. The full CoC will be placed on the platform when it goes live.

#### 3.1.3. Community engagement

In order to measure community engagement directly during the testing and validation phase of the platform, the so-called ‘Voice of the Customer’ approach was/is utilized. This is one of the most popular Agile techniques to capture product functionality as well as user needs and connects the PEERS platform directly with those who are likely to engage with it while also taking their valuable feedback on board. We aim to do this in various ways: (i) obtain user feedback following the product launch via succinct surveys and offer them an attractive opportunity to beta-test new protocols prior to release on the platform; (ii) interview users about their research problems, how they address them and how PEERS could aid their requirements; (iii) listen to users and implement new features and functionalities to the PEERS platform. This approach will help to identify the most vital protocols and facilitate the development of PEERS together with the user and maximize its usefulness. Initial feedback from end user interviews and a small survey suggests a willingness not only to use the PEERS platform to search for information but indeed also to act as a contributor to complete and update any relevant protocols.

Further, PEERS will establish its presence via social media websites (e.g. Researchgate, Twitter, LinkedIn and others) to update the scientific community regularly on new developments and to recruit reviewers for newly added protocols via ‘call-to-action’ announcements. Other indirect metrics to ensure the uptake and adoption of PEERS by the wider scientific community and measure success will employ popular mechanisms utilized by online publications such as the number of hits / user access for each protocol and factor, the number of downloads of the evidence related to each factor and the number of downloads for the citated publications. Information will be graphically displayed on the platform and updated instantly. We can use this approach to identify a) popular protocols; b) protocols with low engagement; c) popular modes of engagement with the different protocols. Outcomes will identify areas of global interest for researchers who use the platform. Simultaneously, this information enables us to identify specific knowledge gaps and we will seek to close them.

### 3.2. Long-term vision of full PEERS

At present, given the short period of its existence, contents of the database is limited, but PEERS aims to upscale and expand by constantly adding new protocols. Given the composition of the Working Group, the list of protocols will be expanded to include other commonly employed, as well as newly developed neuroscience methods such as *in vivo/in vitro* electrophysiology, cell culture (2D+3D), optogenetics, elevated plus maze, qPCR, flow cytometry, light-dark box, conditioned fear etc. The database will continue to be curated and updated for already published protocols.

In the longer term, PEERS aims to attract a broader user base and therefore, the ambition is to branch out and include protocols from other biological disciplines such as infection, inflammation, immunity, cardiovascular sciences, microbial research, etc. Furthermore, the inclusion of expert unpublished data and information related to the importance of specific factors may also be warranted. However, strict rules would need to be set out to ensure proper management, quality control and utility of such unpublished data.

As one of the next steps to aid this expansion, we seek to establish a ‘Board of Editors’ of PEERS akin to an editorial board of a scientific journal, in which all biological disciplines will be represented. Novel protocols can be commissioned accordingly, and the wiki-like structure of the platform would then persist with the Board of Editors reviewing contributions to ensure the extraction of evidence for specific factors is appropriate. The members of the Board of Editors alongside the contributors will be displayed on the platform to make everyone’s contribution transparent.

Most importantly, as PEERS does not compete with existing initiatives for the reporting of results or with guidelines for scientific conduct, we envisage interactions with initiatives such as ARRIVE, EQIPD, FAIR and others to be fruitful in increasing the quality and reproducibility of research in the future.

## Author Contributions

All members of the PEERS Working Group were involved in the preparation of this manuscript (with AS, CD, CFB, AH, MJK, NK, GR, CHE responsible for the scientific aspects (‘scientific arm’) and KK and KP responsible for the technical aspects (‘software arm’). AS, GR and CHE were responsible for the main text of the manuscript, while CHE was responsible for the production of the figures. All other authors (AB, MJP, PP, TS) provided valuable input to early PEERS concepts and edits to the manuscript. All authors reviewed the final version of the manuscript.

## Acknowledgements

We would like to thank IJsbrand Jan Aalbersberg, Natasja de Bruin, Philippe Chamiot-Clerc, Anja Gilis, Lieve Heylen, Martine Hofmann, Patricia Kabitzke, Isabel Lefevre, Janko Samardzic, Susanne Schiffmann and Guido Steiner for their valuable input and discussions during the conceptualization of PEERS and the initial phase of the project.

## Supplementary Information

**Table S1:**
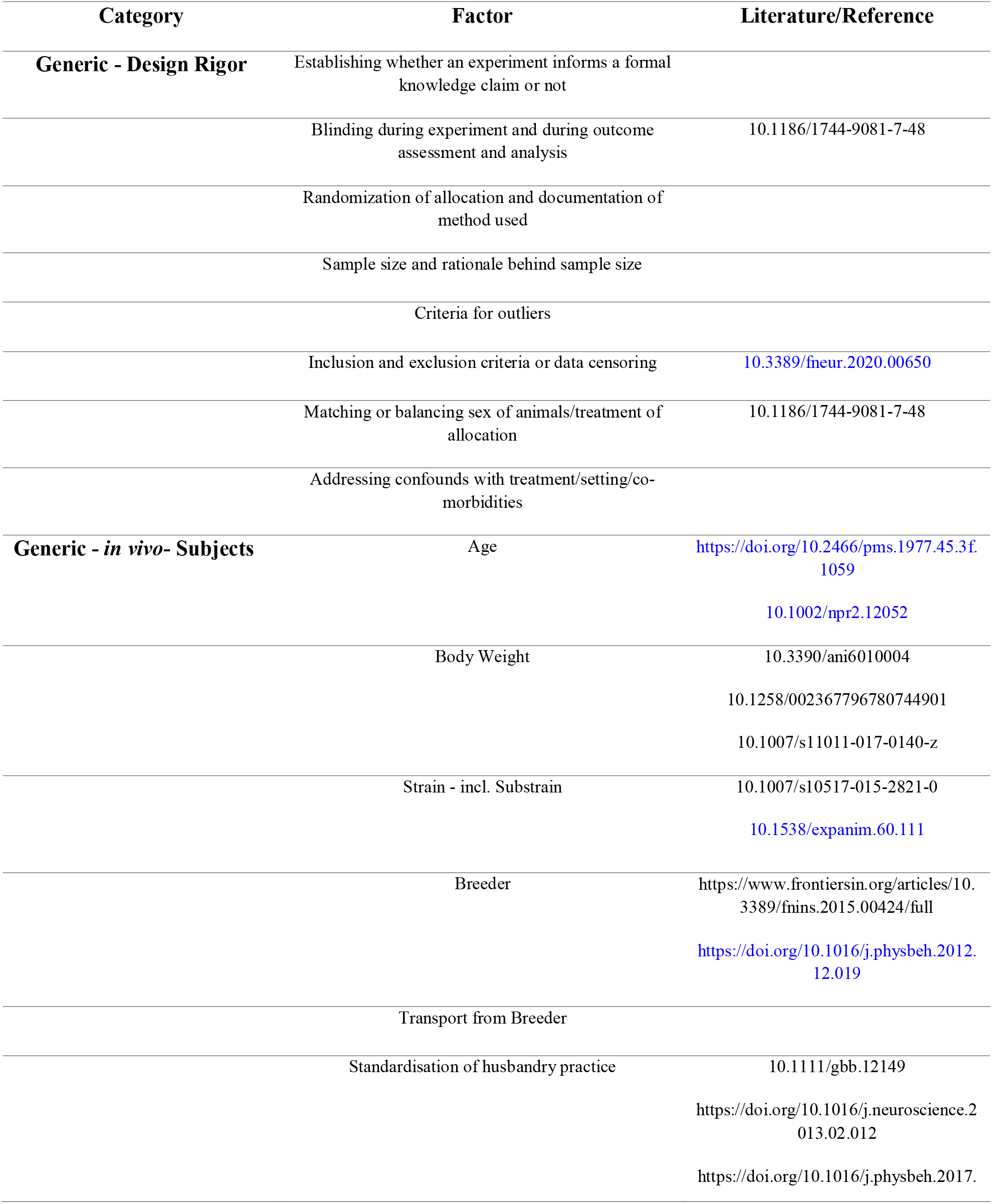
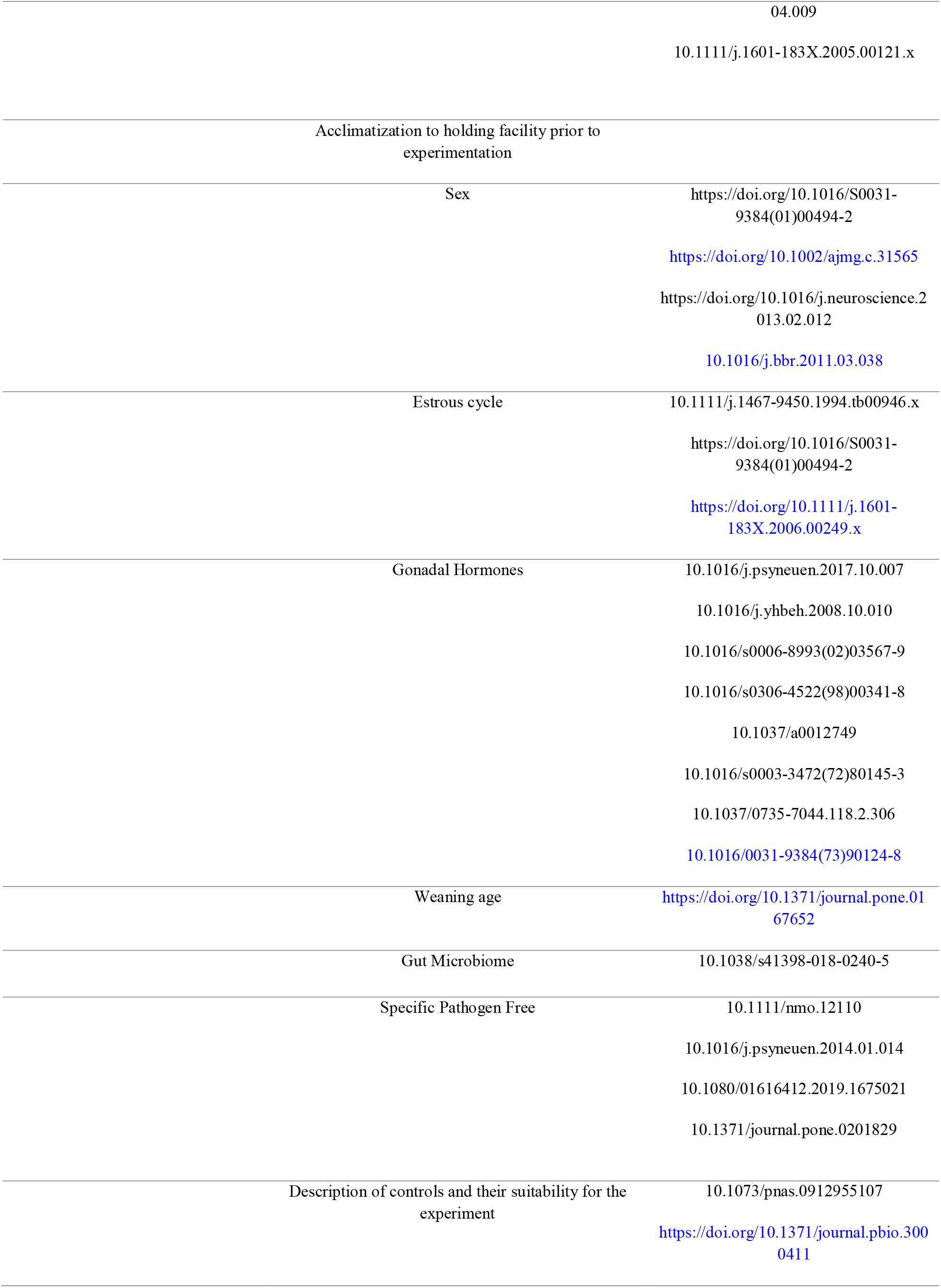
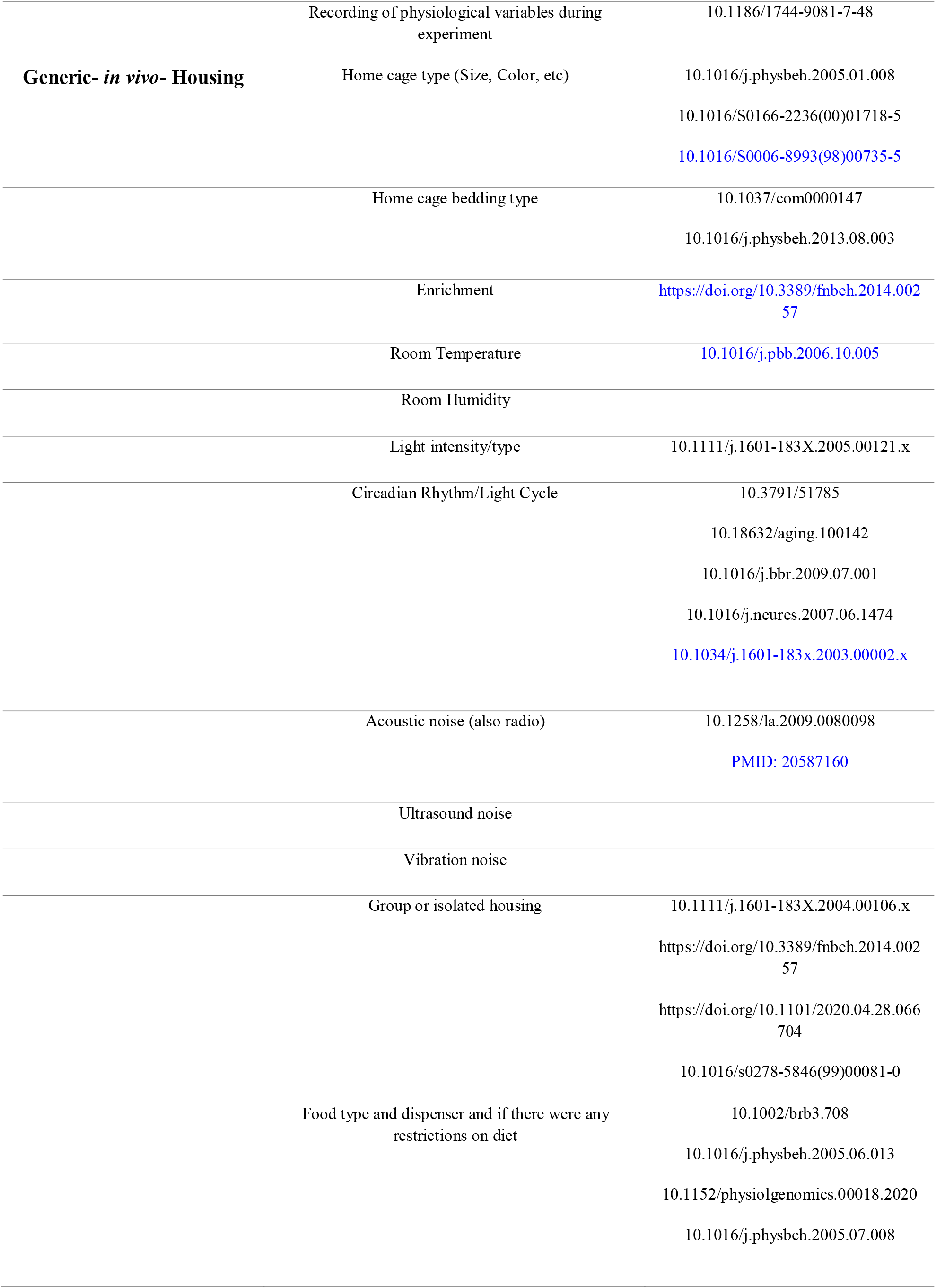
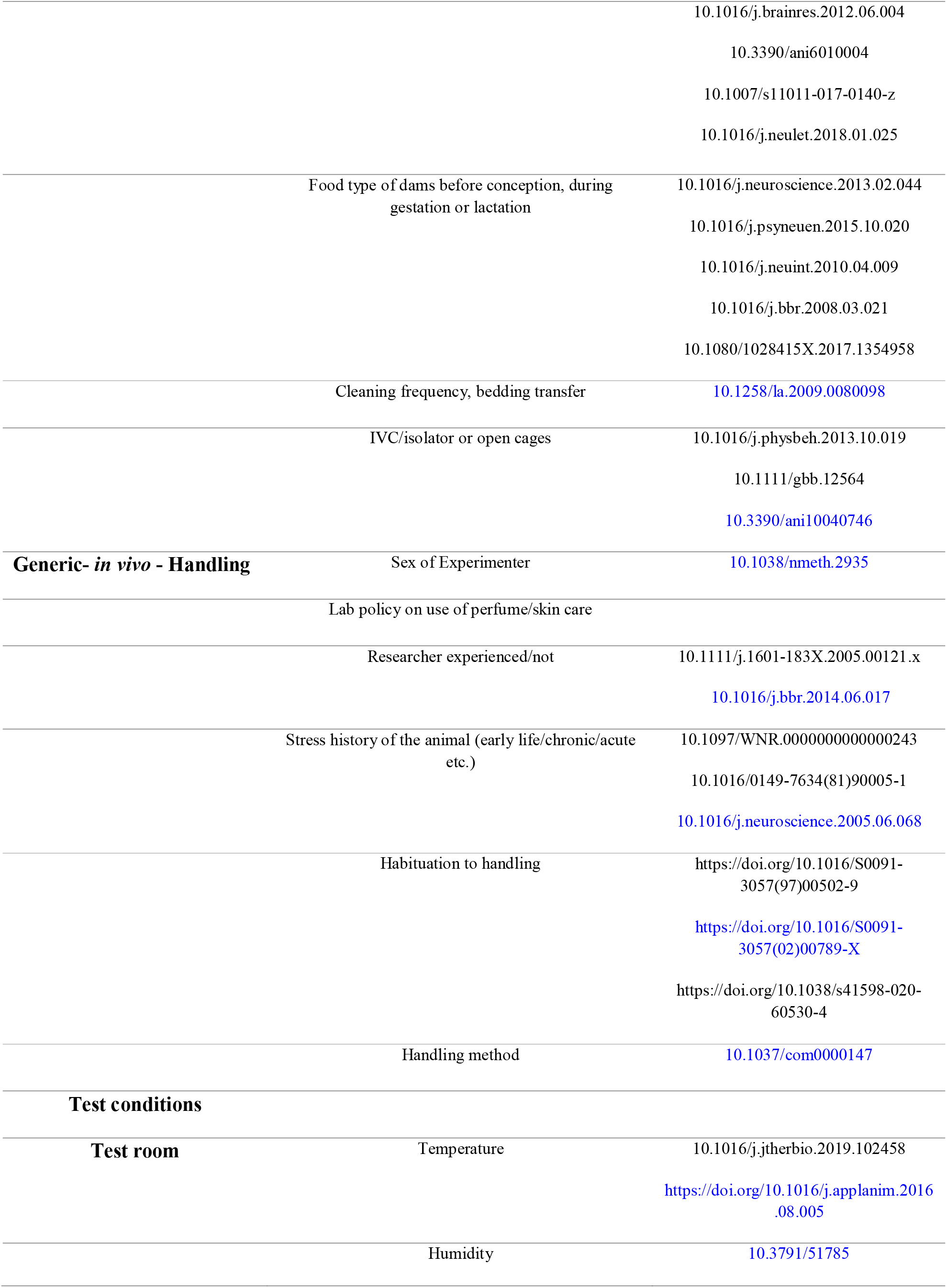
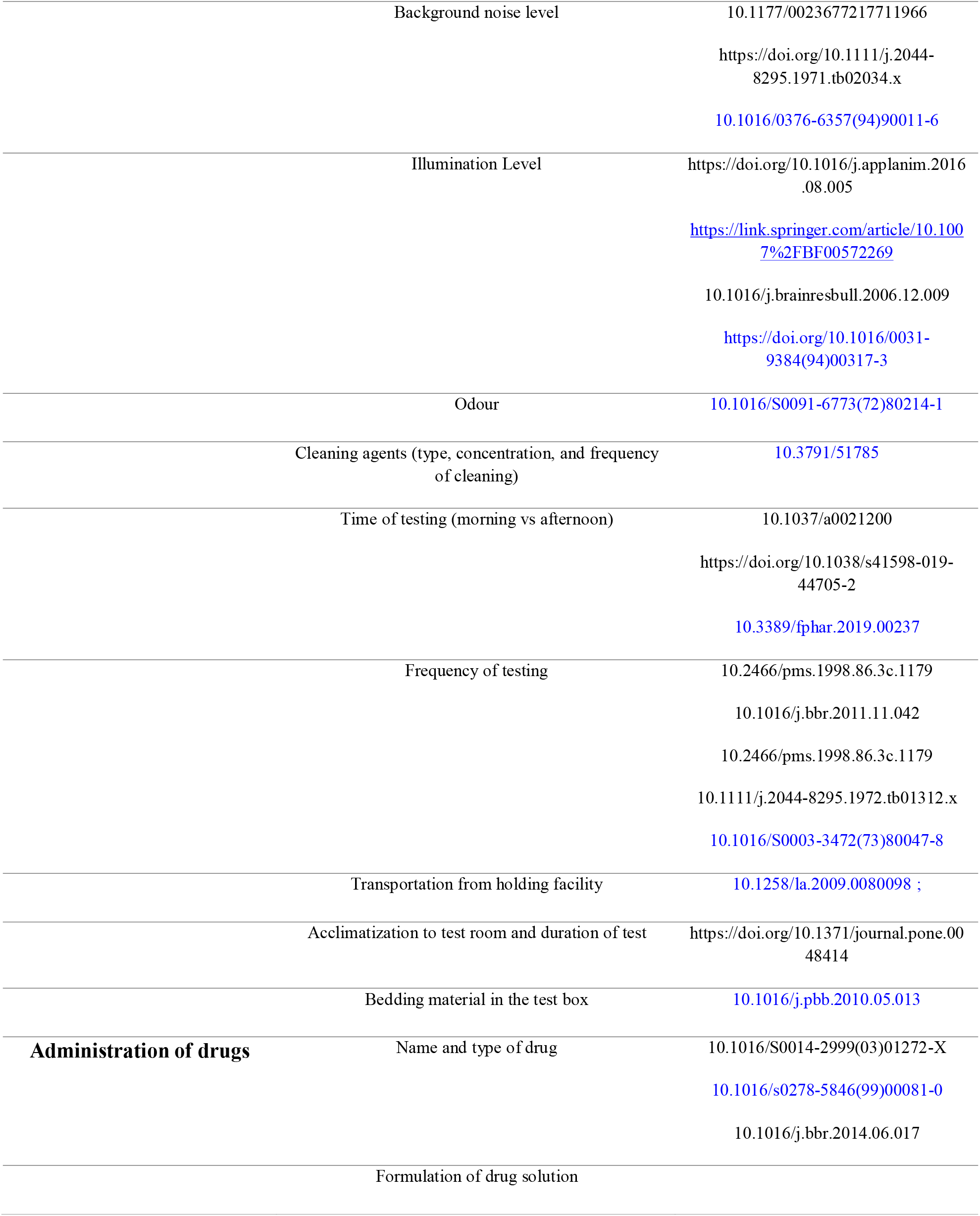
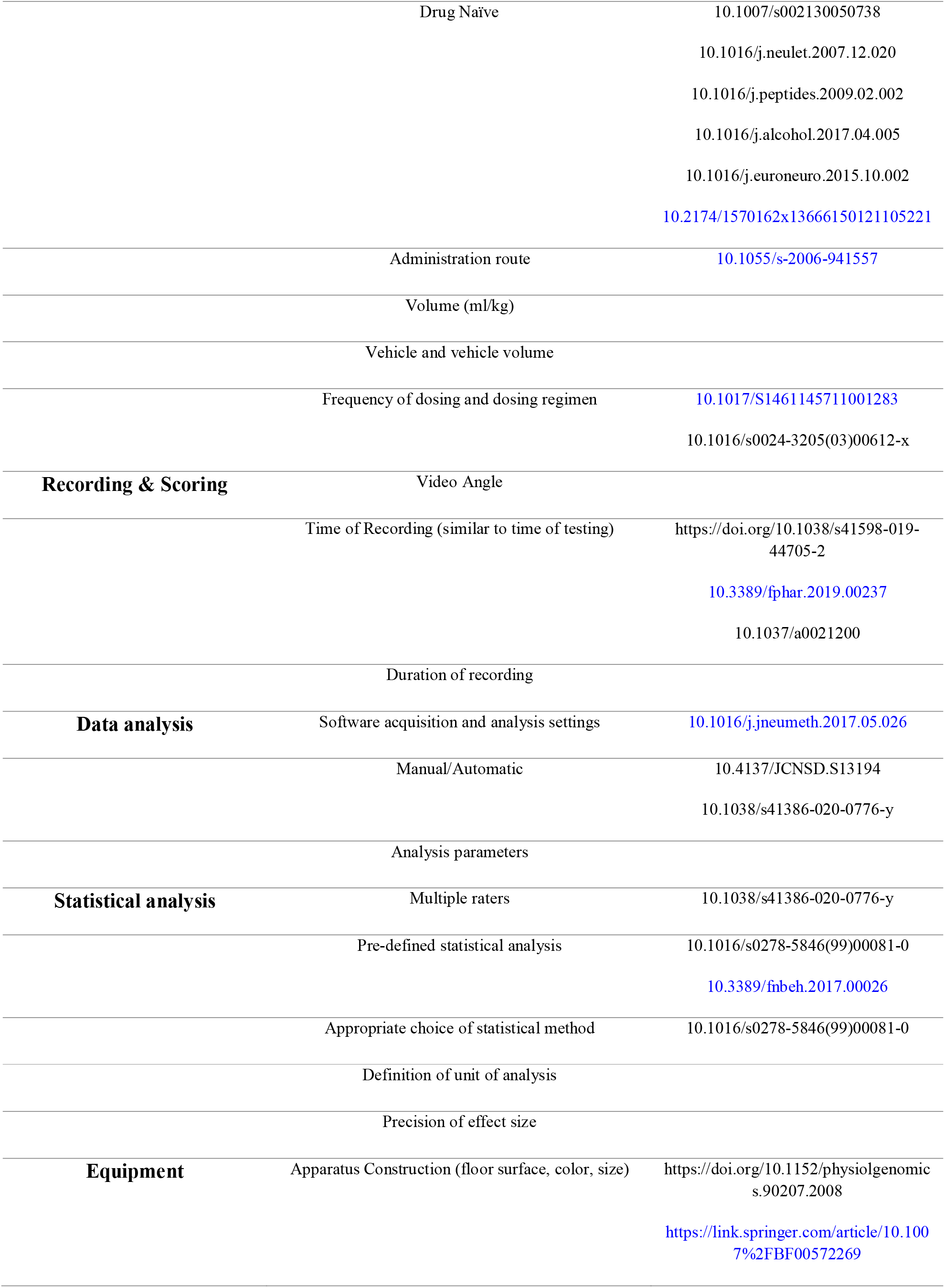
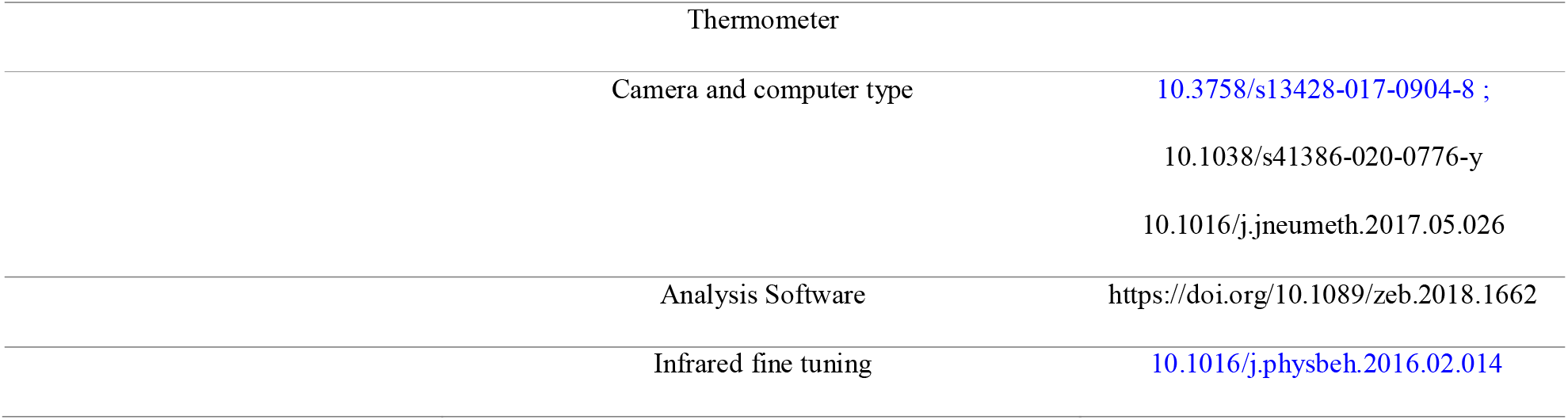
Table of factors with references embedded for the Open Field protocol.

**Table S2:**
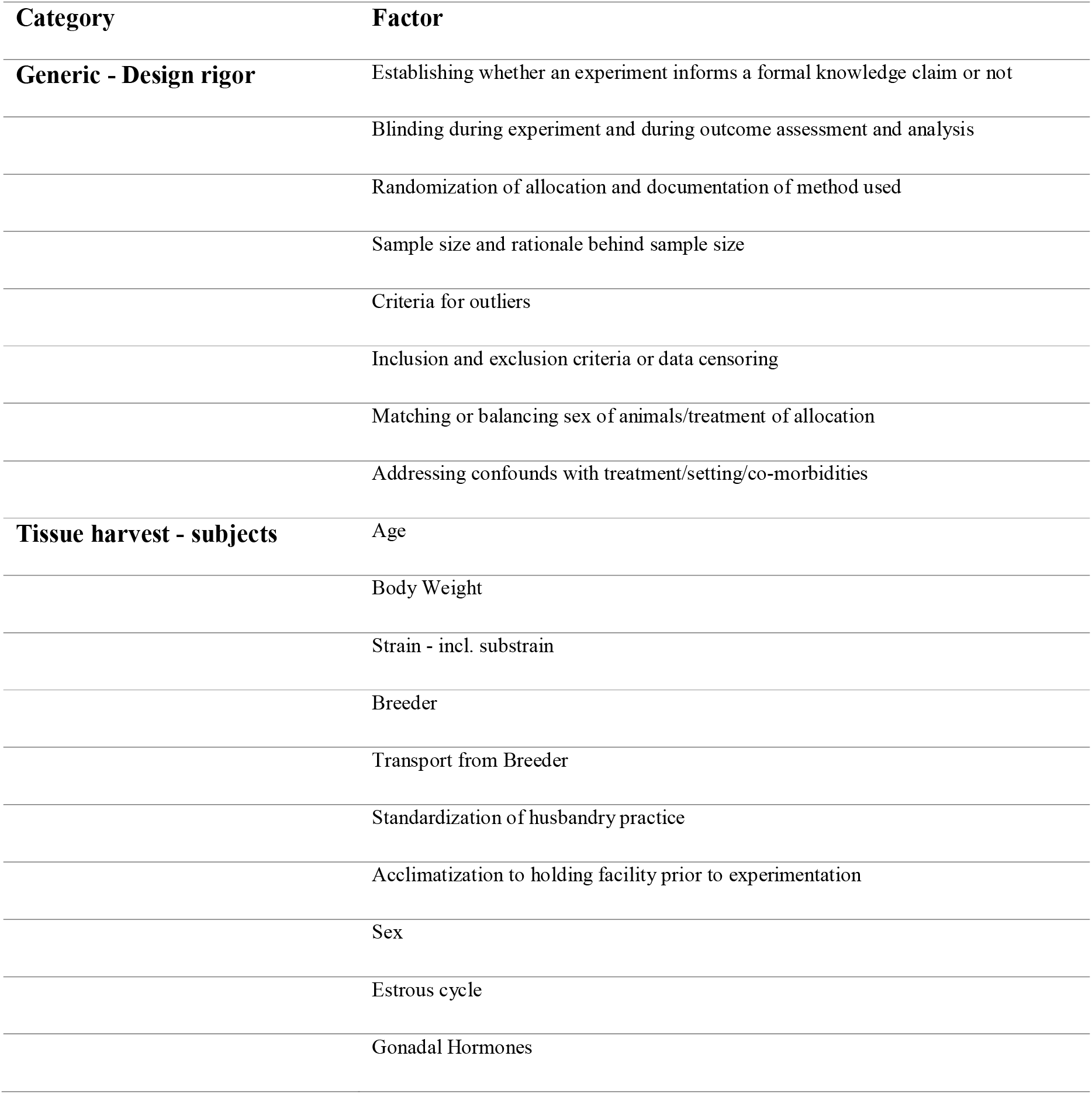
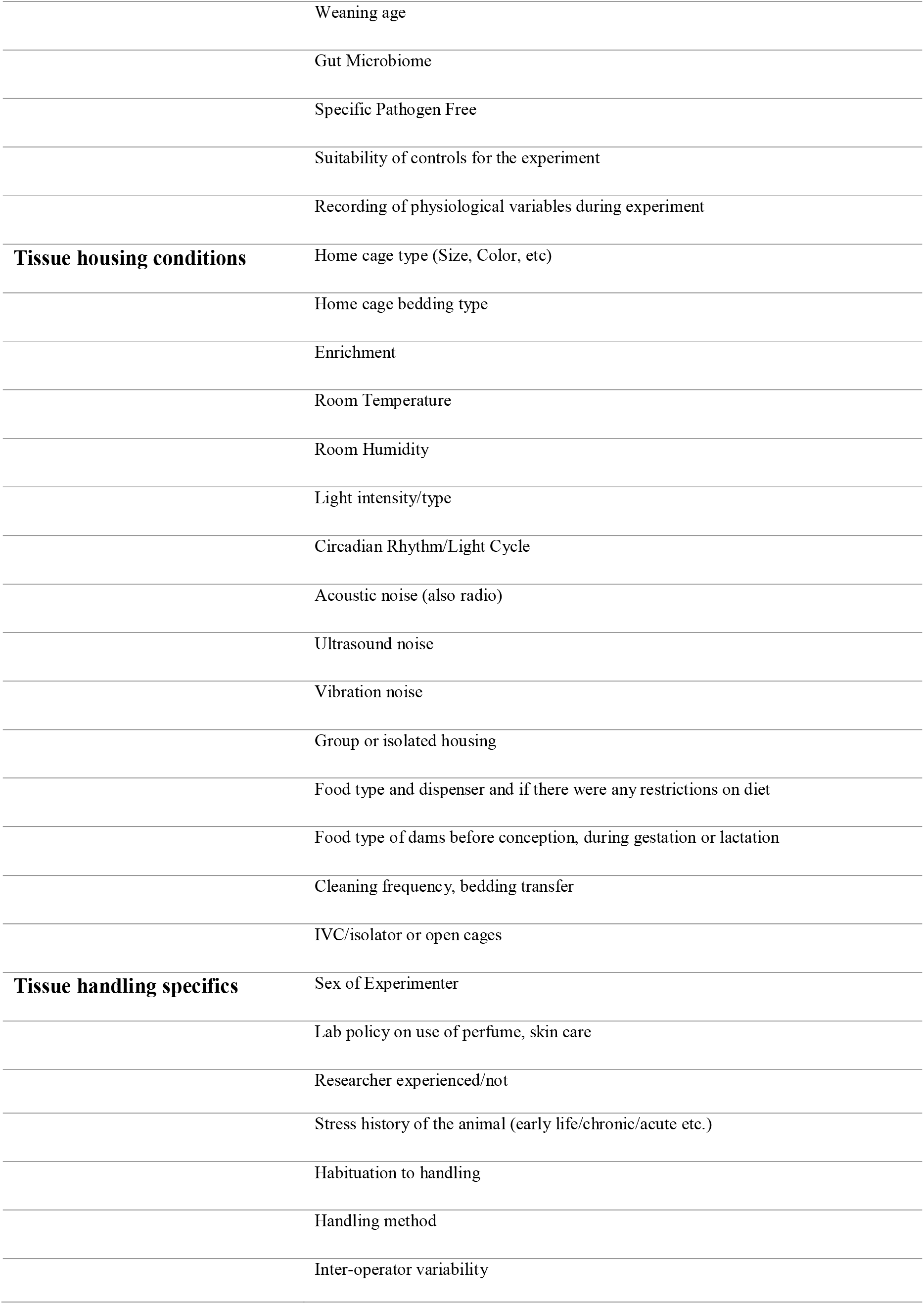
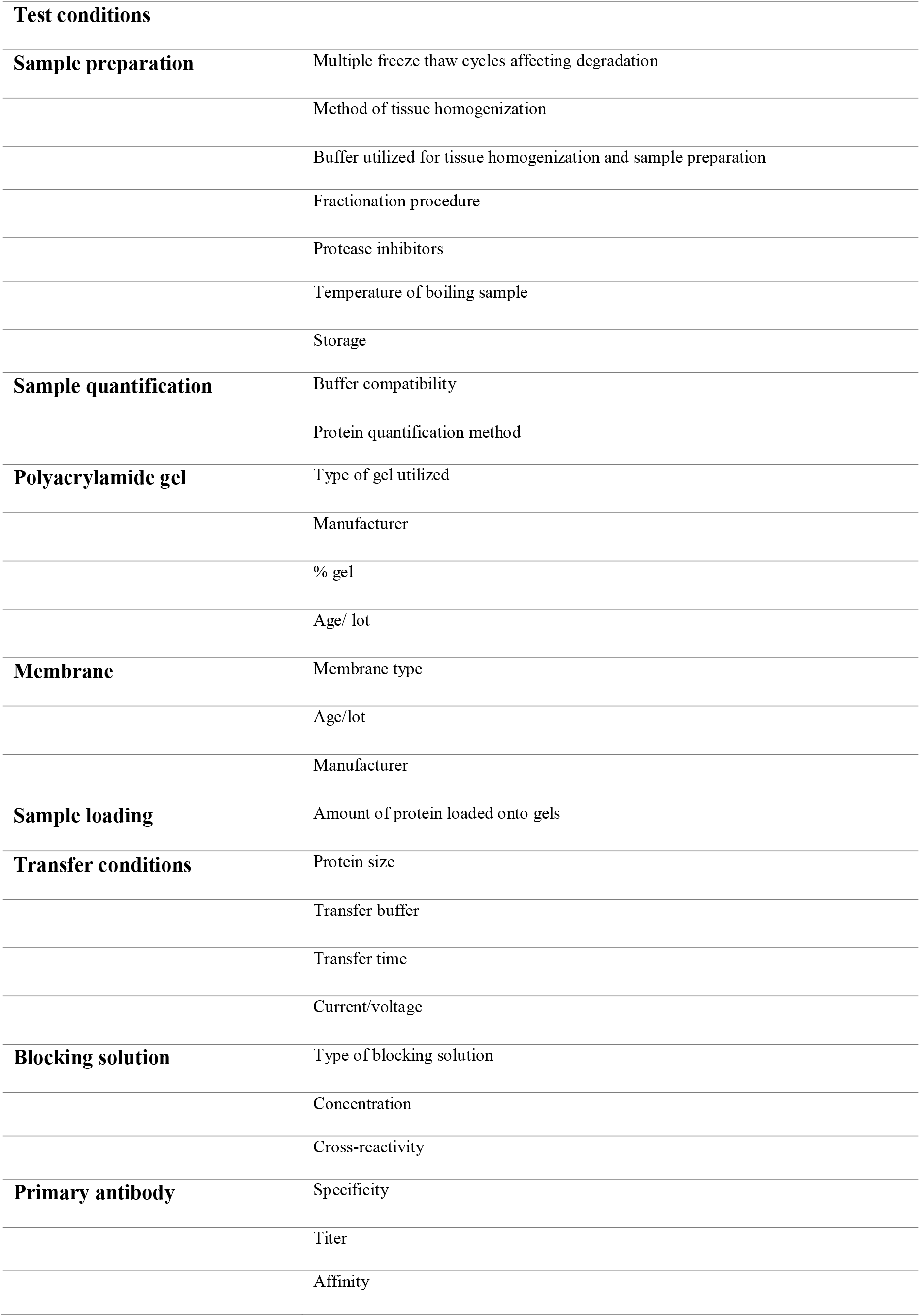
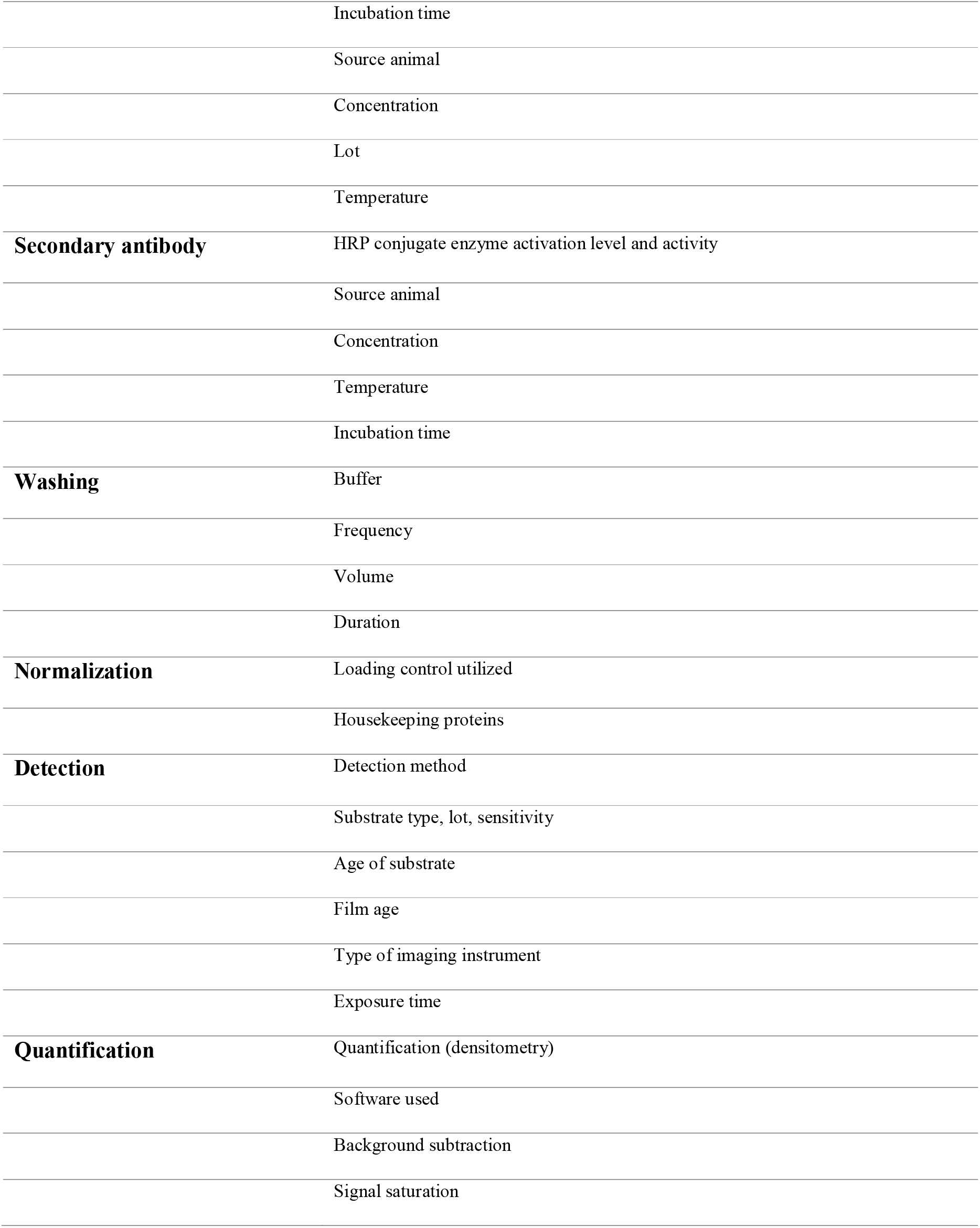
List of factors for the Western Blotting protocol.

| Category | Factor |
| --- | --- |
| **Generic - Design rigor** | Establishing whether an experiment informs a formal knowledge claim or not |
|  | Blinding during experiment and during outcome assessment and analysis |
|  | Randomization of allocation and documentation of method used |
|  | Sample size and rationale behind sample size |
|  | Criteria for outliers |
|  | Inclusion and exclusion criteria or data censoring |
|  | Matching or balancing sex of animals/treatment of allocation |
|  | Addressing confounds with treatment/setting/co-morbidities |
| **Tissue harvest - subjects** | Age |
|  | Body Weight |
|  | Strain - incl. substrain |
|  | Breeder |
|  | Transport from Breeder |
|  | Standardization of husbandry practice |
|  | Acclimatization to holding facility prior to experimentation |
|  | Sex |
|  | Estrous cycle |
|  | Gonadal Hormones |
|  | Weaning age |
|  | Gut Microbiome |
|  | Specific Pathogen Free |
|  | Suitability of controls for the experiment |
|  | Recording of physiological variables during experiment |
| **Tissue housing conditions** | Home cage type (Size, Color, etc) |
|  | Home cage bedding type |
|  | Enrichment |
|  | Room Temperature |
|  | Room Humidity |
|  | Light intensity/type |
|  | Circadian Rhythm/Light Cycle |
|  | Acoustic noise (also radio) |
|  | Ultrasound noise |
|  | Vibration noise |
|  | Group or isolated housing |
|  | Food type and dispenser and if there were any restrictions on diet |
|  | Food type of dams before conception, during gestation or lactation |
|  | Cleaning frequency, bedding transfer |
|  | IVC/isolator or open cages |
| **Tissue handling specifics** | Sex of Experimenter |
|  | Lab policy on use of perfume, skin care |
|  | Researcher experienced/not |
|  | Stress history of the animal (early life/chronic/acute etc.) |
|  | Habituation to handling |
|  | Handling method |
|  | Inter-operator variability |
| **Test conditions** |  |
| **Sample preparation** | Multiple freeze thaw cycles affecting degradation |
|  | Method of tissue homogenization |
|  | Buffer utilized for tissue homogenization and sample preparation |
|  | Fractionation procedure |
|  | Protease inhibitors |
|  | Temperature of boiling sample |
|  | Storage |
| **Sample quantification** | Buffer compatibility |
|  | Protein quantification method |
| **Polyacrylamide gel** | Type of gel utilized |
|  | Manufacturer |
|  | % gel |
|  | Age/ lot |
| **Membrane** | Membrane type |
|  | Age/lot |
|  | Manufacturer |
| **Sample loading** | Amount of protein loaded onto gels |
| **Transfer conditions** | Protein size |
|  | Transfer buffer |
|  | Transfer time |
|  | Current/voltage |
| **Blocking solution** | Type of blocking solution |
|  | Concentration |
|  | Cross-reactivity |
| **Primary antibody** | Specificity |
|  | Titer |
|  | Affinity |
|  | Incubation time |
|  | Source animal |
|  | Concentration |
|  | Lot |
|  | Temperature |
| **Secondary antibody** | HRP conjugate enzyme activation level and activity |
|  | Source animal |
|  | Concentration |
|  | Temperature |
|  | Incubation time |
| **Washing** | Buffer |
|  | Frequency |
|  | Volume |
|  | Duration |
| **Normalization** | Loading control utilized |
|  | Housekeeping proteins |
| **Detection** | Detection method |
|  | Substrate type, lot, sensitivity |
|  | Age of substrate |
|  | Film age |
|  | Type of imaging instrument |
|  | Exposure time |
| **Quantification** | Quantification (densitometry) |
|  | Software used |
|  | Background subtraction |
|  | Signal saturation |

